# Pan-genome analysis reveals the molecular basis of niche adaptation of *Staphylococcus epidermidis* strains

**DOI:** 10.1101/604629

**Authors:** Fei Su, Rui Tian, Yi Yang, Lihui Zou, Xiaomao Xu, Dongke Chen, Junhua Zhang, Xue Chen, Fei Xiao, Gang Zhao, Yanming Li, Hongtao Xu

## Abstract

*Staphylococcus epidermidis* is the most commonly isolated species from human skin and the second leading cause of bloodstream infections. Here, we performed a large-scale comparative study without any pre-assigned reference to identify genomic determinants associated with their diversity and adaptation as a “double-side spy”, a skin dominant colonization, and a successful pathogen. The pan-genome of *S. epidermidis* is open with 435 core proteins and a pan-genome size of 8034 proteins. Genome-wide phylogenetic tree shows that whole genome sequence is a powerful tool to analyze the complex evolutionary process of *S. epidermidis* and investigate the source of infection. Comparative genome analyses demonstrate the high diversity of antimicrobial resistances, especially mobile genetic elements. The complicated relationships of host-bacterium and bacterium-bacterium help *S. epidermidis* to play a vital role in balancing the epithelial microflora. The highly variable and dynamic nature of the *S. epidermidis* genome may be the result of its success in adapting to broad habitats, which is necessary to deal with complex environments. This study gives the general landscape of *S. epidermidis* pan-genome and provides valuable insights into mechanisms for genome evolution and lifestyle adaptation of this ecologically flexible species.

## Introduction

The coagulase-negative *Staphylococcus epidermidis* (*S. epidermidis*) is a common human skin commensal bacterium that can be cultured from virtually every body surface of healthy individuals. It also plays a central role in the skin microbiome [1, 2], it can keep the ecological balance of human skin microflora [3]. *S. epidermidis* can produce various bacteriocins, which kill other microorganisms and have frequently been proposed to enhance survival of the producer strains in a competitive fashion [4, 5]. Especially, serine protease Esp, secreted by *S. epidermidis*, can inhibit the biofilm formation of *S. aureus* and destroy pre-existing *S. aureus* biofilms [6].

However, *S. epidermidis* is the second most common cause of nosocomial infections, which in most cases are antibiotic-resistant [1, 7]. Antibiotic resistance remarkably complicates the treatment and increases the medical expenses [8]. The large gene pool of antibiotic resistance in *S. epidermidis* is shared with many other pathogenic species such as *S. aureus* [9]. Mobile genetic element, multidrug-resistant conjugative plasmids, arginine catabolic mobile element (ACME) [9], and staphylococcal chromosome cassette *mec* (SCC*mec*) elements [10] conferring β-lactam resistance are transferred frequently, enabling rapid evolution and adaptation against antibiotic selection pressure [11, 12]. When the protective layer of the human epithelium is breached and the mechanisms of host immunity fail, staphylococcal infections can become extremely dangerous and even fatal [13]. *S. epidermidis* is particularly associated with the increased use of indwelling medical devices such as artificial heart valves, prosthetic joints, and vascular catheters, which provide a substrate for biofilm formation. On the other hand, during the long-time “arms race”, human beings have developed versatile immunity system with antimicrobial peptides (AMPs) as the first line of innate immune defense on the human skin; meanwhile, *S. epidermidis* also owns multiple mechanisms such as surface charge alteration, extracellular proteases, exopolymers, and efflux pump proteins to fight against AMPs [7]. The complex host-bacterium and bacterium-bacterium relationships make it necessary to investigate the genetic diversity, genome evolution, and lifestyle adaptation of *S. epidermidis*.

Much attention has been focused on understanding the evolution and spread of *S. epidermidis* by different methods [11, 14]. As the time goes on, high throughput sequencing is now fast and cheap and a large amount of genomics data about *S. epidermidis* are accumulated, it is essential to perform more comprehensive comparative and evolutionary study of ecologically diverse strains of *S. epidermidis* for better clinical management. Here, we compared the genomic features of *S. epidermidis* isolates of clinical and non-clinical relevance by using a pan-genome analysis of 198 publicly available *S. epidermidis* strains at the GenBank database of National Center for Biotechnology Information at April 30 of 2017. We assembled the consensus “pan-chromosome” without any pre-assigned genome reference and identified both core and variable regions within the chromosome. Second, we utilized a comparative genomics approach on 198 genomes to analyze the diversity of antibiotic resistance of *S. epidermidis*. Our results revealed that *S. epidermidis* isolates encoded a vast collection of genetic determinants and mechanisms to confer antibiotic resistance, antimicrobial peptides resistance, and survival adaptations. These analyses will provide insight into the coevolution of *S. epidermidis* as a nosocomial pathogen and directly aid the future efforts for large-scale epidemiological studies of this continuously evolving multi-drug resistant organism.

## Methods

### Strains

A total of 198 *S. epidermidis* isolates were selected to represent known diversity within the species and multiple locations and sources until Apirl 30 of 2017, including reference genomes from S. epidermidis strain RP62A (Gill et al. 2005). All the available genome sequence of *S. epidermidis* strains and related annotation data were downloaded through the GenBank database [15] of NCBI (see Table S1 in the supplemental material).

### SCC*mec* and ACME typing

An SCC*mec* sequence cassette database was prepared with the following accession numbers downloaded from NCBI: AB033763.2 (Type I), AB433542.1 (Type I.2), D86934.2 (Type II), AB261975.1 (Type II.4), AJ810123 (Type II-B), AB127982.1 (Type II-B), AM983545.1 (Type II-D), HE858191.1 (Type II-E), AB037671.1 (Type III), HM030721.1 (Type IV), HM030720.1 (Type IV), AM292304.1 (ZH47 mobile elements), AB425824.1 (Type IV), EU437549.2 (Type IV-A), AB063172.2 (Type IV-A), AB063173 (Type IV-B), AY271717.1 (Type IV-C), AB096217 (Type IV-C), AB245470.1 (Type IV-C), AB097677.1 (Type IV-D), AJ810121.1 (Type IV-E), DQ106887.1 (Type IV-G), AB633329.1 (Type IV-I), AB425823.1 Type IV), AB121219.1 (Type V), AB478780.1 (Type V), AB512767.1 (Type V), AF411935.3 (Type VI), AB462393.1 (Type VII), AB373032.1 (Type V-C1), FJ670542.1 (Type VIII), FJ390057.1 (Type VIII), AB505628.1 (Type IX), AB505630.1 (Type X), and FR821779.1 (Type XI) [16].

The ACME-*arc*A and ACME-*opp*3AB genes were used as markers of the ACME-*arc* cluster and the ACME-*opp*3 cluster, respectively. ACME was classified as type I (contains the ACME-*arc*A and ACME-*opp*3AB gene clusters), type II (carries only the ACME-*arc*A locus), and type III (carries only the ACME-*opp*3AB locus) [17]. ACME-*arc*A and ACME-*opp*3AB identified in this study were compared with the reference sequences of ACME-*arc*A (USA300_FPR3757) and ACME-*opp*3AB (USA300_FPR3757).

### kSNP *S. epidermidis* trees

A phylogenetic tree was inferred from single-nucleotide polymorphisms (SNPs) identified by kSNP (version: 3.0, https://sourceforge.net/projects/ksnp/) [18] by using a *k*-mer length 19 nucleotides and based on a requirement that at least 80% of the genomes have a nucleotide at a given SNP position in order for the SNP to be considered to be a core and included in tree building. A total of 1832 core SNP positions were identified. These SNPs were used to infer a maximum-likelihood tree with RAxML [19] with 100 bootstrap replicates.

### Pan-genome analysis

Cluster of orthologous proteins were generated with version 3.24 of PanOCT (https://sourceforge.net/projects/panoct/) as previous described [20]. Briefly, PanOCT deals with recently diverging paralogs by using neighborhood gene information. All the parameters were set to default values except for the length ratio to discard shorter protein fragments when a protein is split due to a frameshift or other mechanisms was set to 1.33 as recommended by the authors. Orthologous clusters were stringently defined as all sequences in a cluster having shared sequence identity ≥ 70 % and coverage ≥ 75 %. Plots and calculations of pan-genome sizes, new genes discovered and pan-genome status were also determined as described previously [21].

### Characterization of strains

*In silico* multilocus sequence typing of 198 strains was performed with the MLST 1.8 online server [22]. The antimicrobial resistance genes in the sequenced isolates were identified by BLASTp [23] searching with the databases of ARDB [24]. Genes conferring virulence factors were identified using BLASTp with VFDB [25]. Given that many virulence factors for *S. epidermidis* that are not contained in the VFDB, we used the orthologous proteins and virulence factors from RP62a [26] and ATCC1228 [27] to make up the missing information.

### Functional analysis

All genes are BLASTed against all sequences in the database of KOBAS 2.0 (http://kobas.cbi.pku.edu.cn/) [28]. The cutoffs are BLASTp *E*-value <10-5 and BLAST subject coverage > 70 %. We used the genes from same genome as the default background distribution and considered only pathways for which there were at least two genes mapped. Fisher’s exact test was choosing to perform statistical test and Bonferroni correction was used to reduce the high overall Type-I error with p.adjust from R package.

### Statistical analyses

The differences in the prevalence of antimicrobial resistance genes and phenotypes among isolates were analyzed by using two-tailed Fisher’s exact test and Bonferroni correction was also performed as mentioned above. All the statistical analyses were carried out using R package (version: 3.3). A *P* value of < 0.05 was regarded as statistically significant.

## Results

### Core pan-genome of *S. epidermidis*

Despite the intensive effort to characterize *S. epidermidis* and the sizable number of whole genome comparisons in literature [29], more and more genome data is rapid accumulated and could easily obtained from public database, such as NCBI. Using PanOCT, a total of 8,034 orthologous protein clusters were identified from a collection of all *S. epidiermidis* genomes publicly available at the time of the analysis (Supplementary Table S1). PanOCT only includes non-paralogs in clusters and uses conserved gene neighborhood to separate duplicated genes. This means that insertion sequence elements that are in novel contexts will often form singleton clusters even though they are identical in sequence to other IS elements within or between genomes analyzed. When the “core” pan-genome is defined to be present at all 198 genomes analyzed, there were 435 (5.4 %) core protein clusters and 2915 (36.3 %) novel clusters (groups with a single member from a single genome) (Fig. 1a). To predict the theoretical maximum pan-genome size (i.e., the total number of genes, including core, unique, and dispensable genes) a pan-genome model was implemented using medians and an exponential decay function (Fig. 1b). The maximum pan-genome size was estimated to be 12,554 ± 65 genes. To determine whether the *S. epidiermidis* pan-genome is open or closed, the number of new genes identified (i.e., unique or strain-specific genes) for each genome added was determined and fit to a power law function (n = κ N^-α^) as described previously [21]. According to the result, we found the pan-genome of *S. epidiermidis* appeared to be open (α = 0.226 ± 0.002; Fig. 1b). For each genome added, the number of new genes was extrapolated by calculating tg(θ), which was determined to be 7.7 ± 0.4 (Fig. 1b).

**Figure 1.**
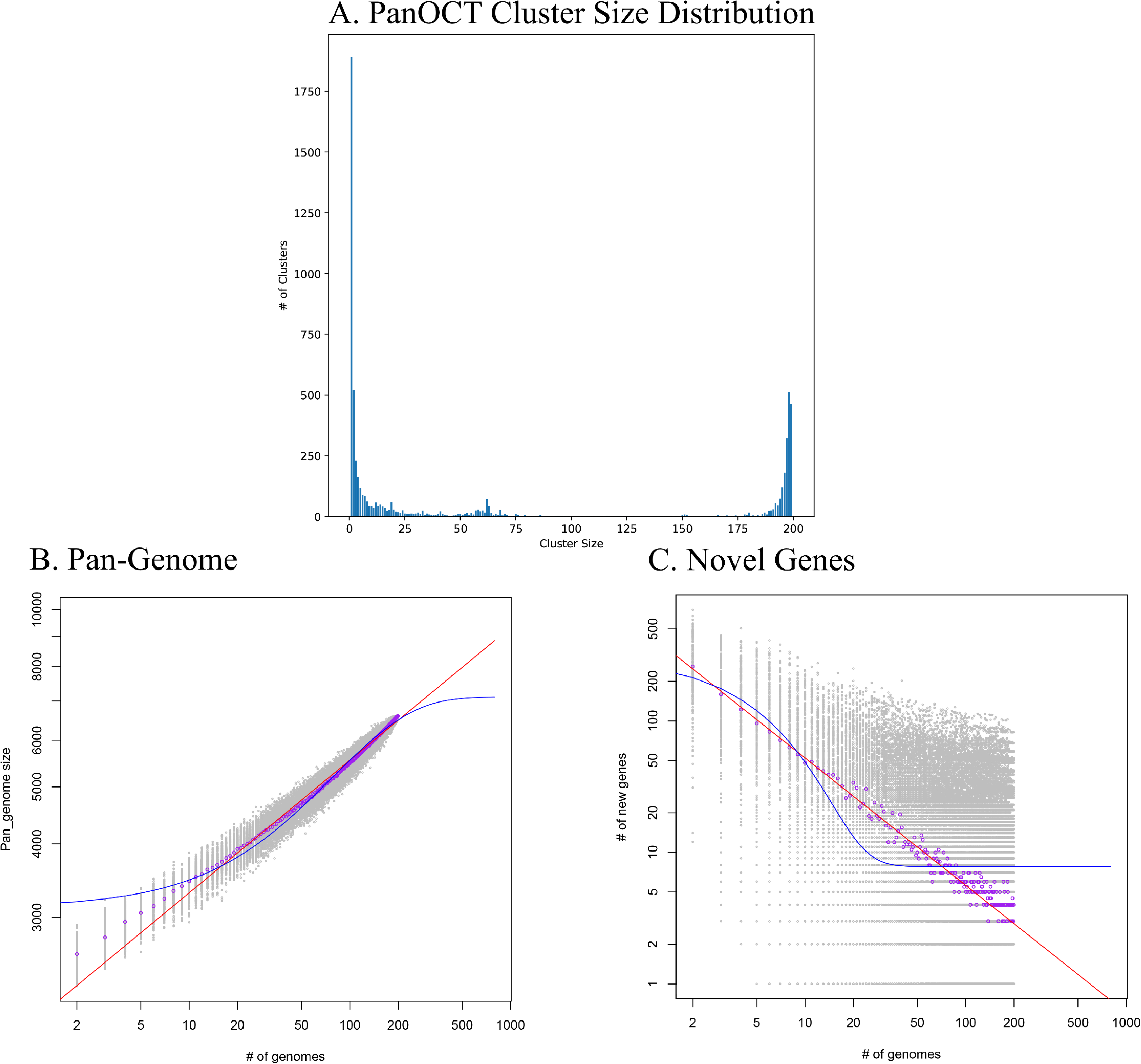
Analysis of the *Staphylococcus epidermidis* pan-genome. (a). The distribution of protein cluster sizes generated from the comparison of 198 *S. epidermidis* genomes using PanOCT. (b). The pan-genome size (left) and the number of novel genes discovered with the addition of each new genome (right) were estimated for all 198 genomes using a pan-genome model based on the original Tettelin et al. model [21].

**Figure 2.**
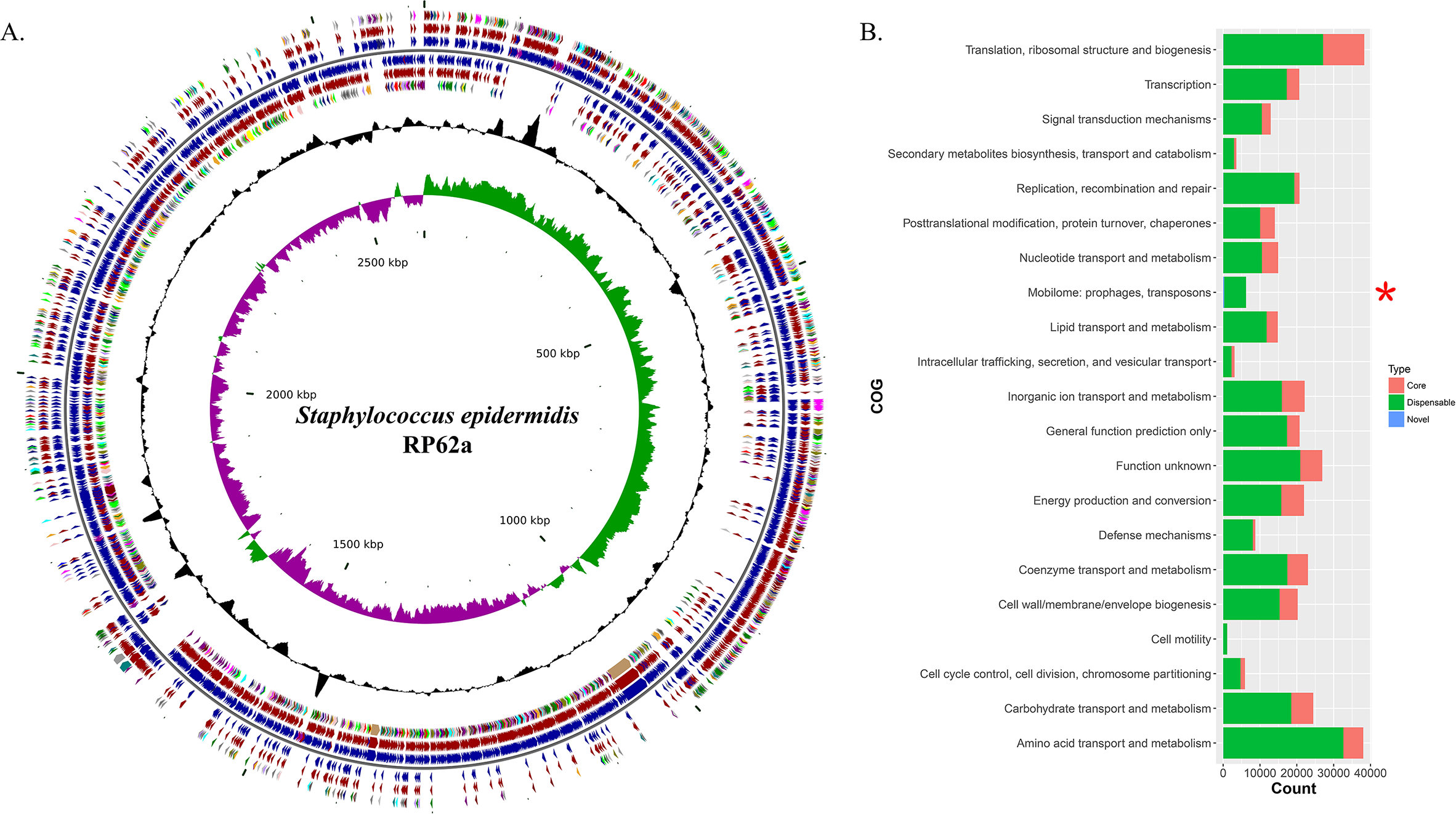
Functional analysis of the pan-genome of *Staphylococcus epidermidis*. (a). Distribution of core / dispensable / novel genes in the type strain RP62a. Starting from the outermost ring the feature rings depict: (1) COG functional categories for forward strand coding sequences; (2) Core (brown) / Dispensable (blue) genes for forward strand coding sequences; (3) Forward strand sequence features; (4) Reverse strand sequence features; (5) Core (brown) / Dispensable (blue) genes for reverse strand coding sequences; (6) COG functional categories for reverse strand coding sequences. (7) GC content; (8) GC skew. The colors of different COG functional categories were following the definition of Grant et al. [46]. (b). Numbers of core, dispensable and novel genes for each COG category. COGs significantly enriched (adjusted *P*-value < 0.05, Fisher exact test) in core, dispensable, or novel genes are marked with red asterisk.

The function of the genes within the variable genome was investigated by assigning all gene clusters to clusters of orthologous groups (COGs) categories [30] and the results showed that novel genes were most likely to be assigned to categories (Supplementary Table S2 and S3) such as mobilome, ribosomal structure and biogenesis, carbohydrate transport and metabolism, and nucleotide transport and metabolism, based on the result of Fisher’s exact test.

### Phylogenetic relationship of *S. epidermidis* isolates

To estimate the genetic relationships among *S. epidermidis* strains, we compared all 198 genomes by using a single nucleotide polymorphism-based phylogeny. SNPs were identified from the combined set of genome sequences by using kSNP. Nucleotide positions present in at least 80 % of all genomes were used to build a Maximum-Likelihood phylogenetic tree with RAxML following the tutorial. Strikingly, the 198 *S. epidermidis* isolates formed two distinct groups (Fig. 3), called Cluster A (solid line) and B (dotted line). Most of Sequence Type (ST) 2 nosocomial isolates were near identical at the nucleotide level for all core genes (Supplementary Table S4). All of ST 2 strains in this study presented in Cluster A and had an extremely short evolutionary distance from each other, indicating that these strains were probably derived from a recent common ancestor. By contrast, Cluster B represents a lineage with reduced virulence and all of ST 5 commensal strains presented in Cluster B and clustered together. The rest of Cluster B had a much longer evolutionary distance from ST 5 strains. This clade may have more complex history of evolution and produce a various sub-group.

**Figure 3.**
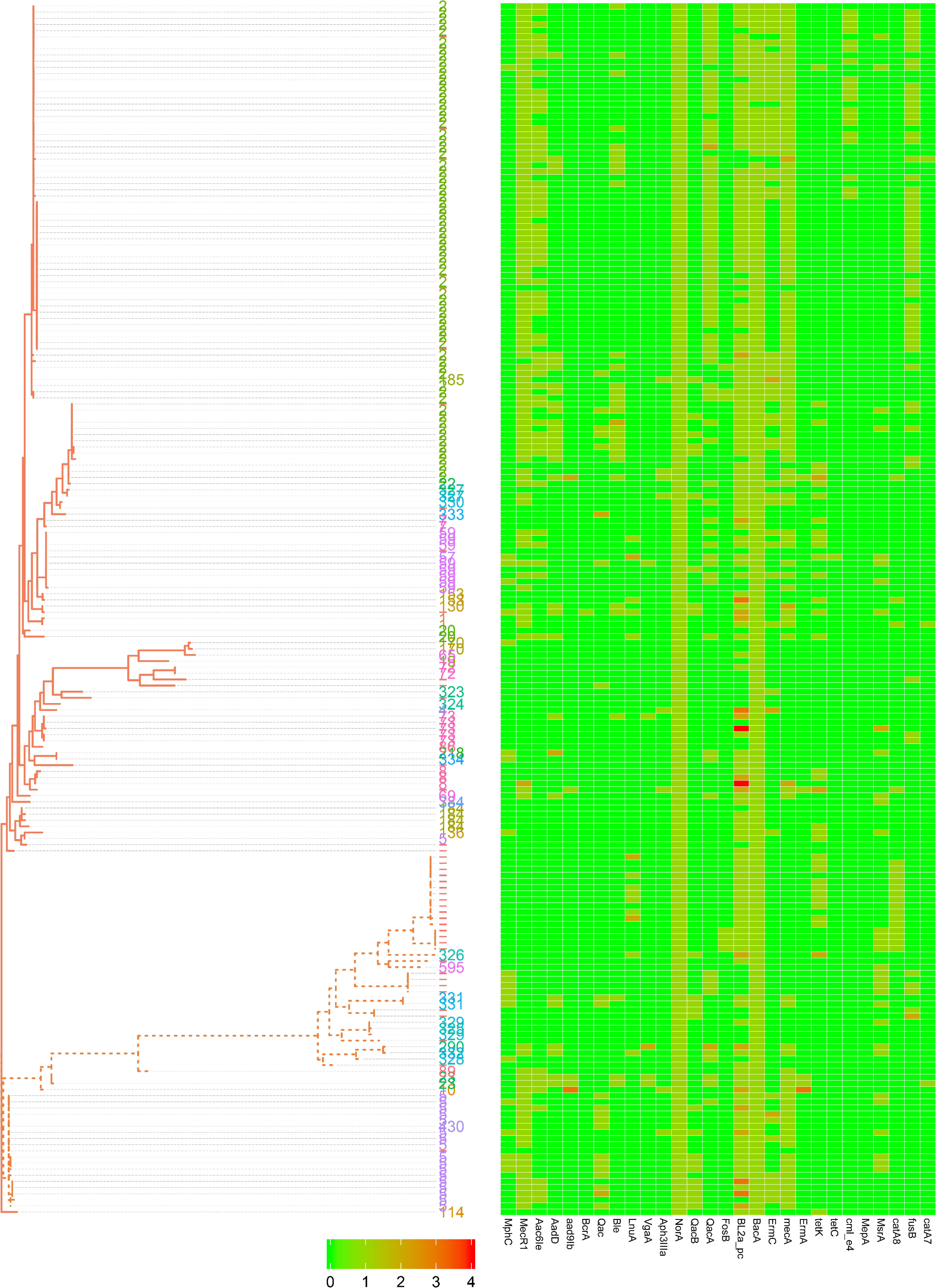
Phylogenetic SNP tree of *Staphylococcus epidermidis* strains. A whole-genome core SNP maximum likelihood tree was constructed for 198 genomes with kSNP and RAxML. Heatmap on the right indicates copies of 28 genes involved in antibiotic resistance. Legends on the bottom stand for copy number of resistant genes.

### Antimicrobial resistance across *S. epidermidis*

Antimicrobial resistance (AMR) is very common among *S. epidermidis* isolates and often limits treatment options [31]. Given the clinical importance of AMR in *S. epidermidis*, we performed a genome-wide analysis of all known AMR genes within our genomic dataset. According to the analysis of ARDB database, we found 28 different types of genes involved in 31 antibiotics (Fig. 3). Nearly all isolates carry at least one type antibiotic resistance gene. Among the genes involved in antimicrobial resistance, our data showed that there were two genes, *nor*A and *bac*A, conserved in all strains. Based on the enrichment analysis of strains from different sources, we found that strains from sources (skin, blood, environment and plant) had significantly different antibiotic resistance profiles: isolates from blood (9 antibiotic resistance genes) and skin (8 antibiotic resistance genes) had significantly enriched antibiotics (Supplementary Table S5), while isolates from environment had no significantly enriched antibiotics. First-line antibiotic therapy for catheter-related bloodstream infections was vancomycin. None of the isolates were resistant to the antibiotic at the genetic level, regardless of isolation source.

### SCC*mec* and ACME in *S. epidermidis*

SCC*mec*, or *staphylococcal* cassette chromosome *mec*, is a mobile genetic element that carries the central determinant for broad-spectrum beta-lactam resistance encoded by the mecA gene a mobile genetic element of *Staphylococcus* bacterial species [10, 32]. According to the completeness of genome in this study (only 7 complete genome sequences), we only analyzed the genes from well-defined SCC*mec* genomic islands [33]. There were 58.6 % (116/198) of *S. epidermidis* strains, in which complete *mec* gene complexes, *mec*A, and *mec*R1 genes were detected (Supplementary Table S1). However, only 39.4 % (78/198) of strains had *ccr* gene complex from type IV cassette, in which both *ccr*A and *ccr*B were present. Similar to the previous results [29, 34], nearly all of the ST2 nosocomial isolates (94.6 %, 70/74) had at least one copy of *mec*A from type IX cassette and *mec*R1 from Type VIII or IV-G cassette. On the other hand, a high prevalence (98 %, 195/198) of ACME was found in *S. epidermidis* strains in this study, of which 22.7 % (45/198) was type I and 75.8% (150/198) was type II.

### Biofilm formation of *S. epidermidis*

Biofilm formation is the major of virulence factor of *S. epidermidis* strains, which will contribute to the persistence of clinical infection. Here, we analyze some well-known genes involved in biofilm formation such as adhesive molecules, including polysaccharide intercellular adhesin (*ica*ABCD), proteinaceous factors (*bhp* and *aap*), teichoic acids, extracellular DNA and so on (Supporting information Table S6). The polysaccharide intercellular adhesion (*ica*ABCD) genes that encode biofilm-associated genes for poly-N-acetylglucosamine synthesis were found in 60% of the commensal isolates, in agreement with previous studies [34]. Especially, any of the *ica* genes was not found in some ST 2 strains (Fig. 4). Gene *aap* was enriched in the blood (adjusted *P*-value < 0.01) compared to the remaining isolates and therefore might be a potential biomarker for *S. epidermidis* infection. We analyze the enrichment of all genes involved in virulence factors and found the *ica*ABCD was significantly enriched despite the sources or sequence types.

**Figure 4.**
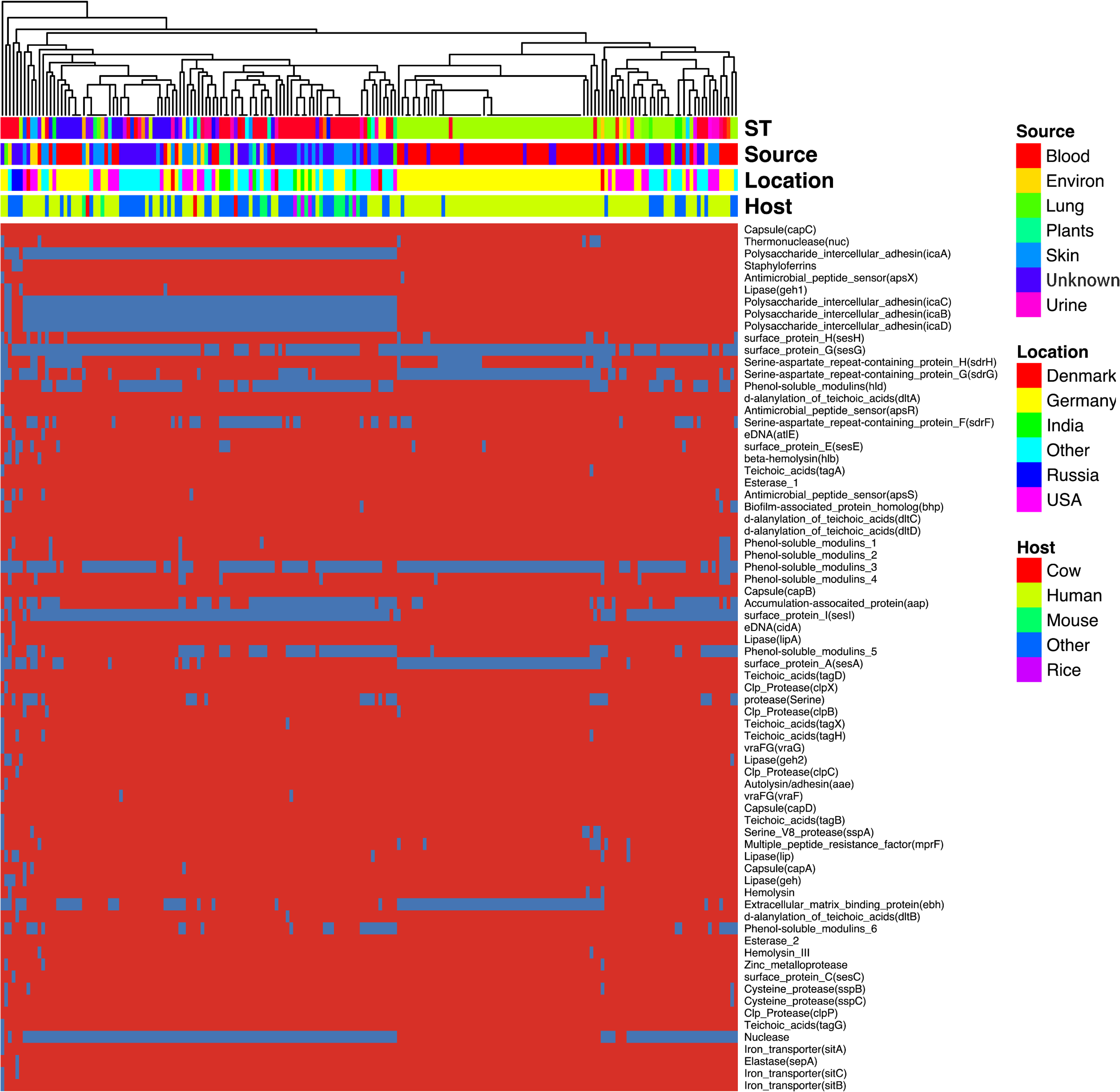
Heatmap of virulence factors among the *Staphylococcus epidermidis* strains. The dendrogram was generated using complete linkage clustering of copies of genes involved in virulence factors. The red color stands for genes that exist in the genomes and the blue color for missing ones. Legends on the right stand for colors of different host, isolates and geographic information.

**Figure 5.**
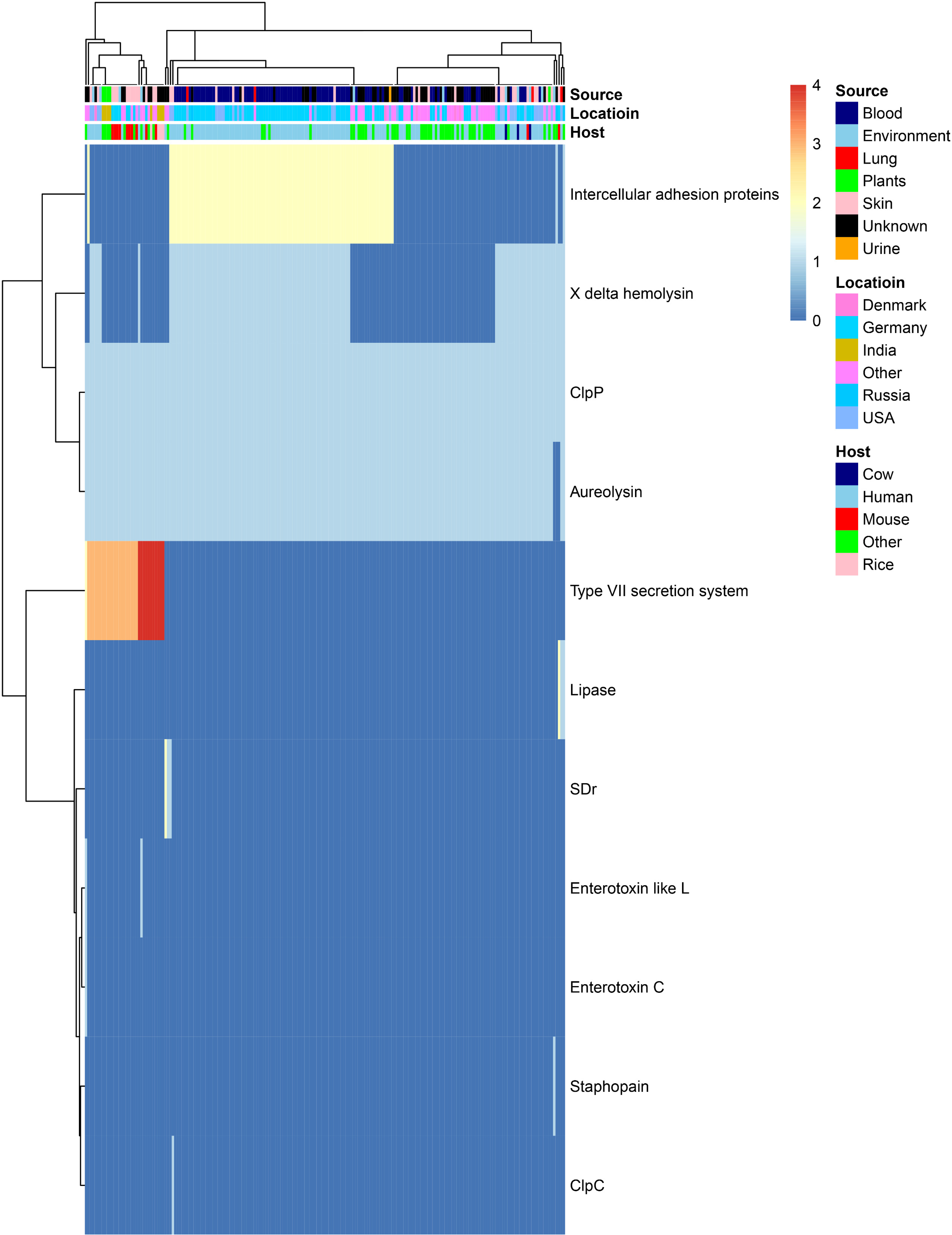
*In silico* analysis of virulence factors of the *Staphylococcus epidermidis* strains. The types of virulence factors were following the VFDBs database. Legends on the right stand for colors of different host, isolates, and geographic information. Different colors stand for copy number of each virulence factors.

### Human-Bacterium and Bacterium-bacterium interactions in *S. epidermidis*

*S. epidermidis* is the major colonization microorganisms in the human skin with complex human-bacterium and bacterium-bacterium interactions. We analyzed the genes (Table 1 and supplementary Table S5) involved in resistance against antimicrobial peptides that can inhibit the growth of most skin microorganism including *S. epidermidis*. Some genes (e.g. *cap*ABCD), which are significantly enriched in the blood and skin, were reported to assist the strain to survive on the skin surface [7]. We also analyzed genes involved in bacterium-bacterium interactions. We found that the genes involved in short-chain fatty acids biosynthesis and extracellular proteases (e.g. Esp) had no difference despite the isolates.

**Table 1.** *S. epidermidis* resistance mechanisms that target AMPs.

| Resistance mechanism | Gene | Target AMPs | Functions | Enrichment |
| --- | --- | --- | --- | --- |
| AMP sensing | *apsSRX* | Most cationic AMPs | 3-component sensor/regulator | - |
|  | *braSR/braDE/vraDE* | Bacitracin, nisin |  | - |
| Phosphatidylglycerol lysylation | *mprF* | Most cationic AMPs | Lysylation of membrane phospholipids | - |
| Teichoic acids alanylation | *dltABCD* | Most cationic AMPs | Alanylation of teichoic acids | Blood / Skin (*dltD*) |
| Exopolymers | *icaADBC* | HBD3, LL-37, DCD-1 | Production of PNAG exopolysaccharide; IcaB<br>*N* -acetylglucosamine<br>deacetylase introduces<br>positive charge | Blood(*icaB*) |
|  | *cap*ABCD | HBD3, LL-37, DCD-1 |  |  |
| Extracellular proteases | *sepA* | LL-37 | Degrades AMPs | - |
|  | *esp* | LL-37a |  | - |
| ABC transporters | *vra*FG | Vancomycin, polymyxin B,<br>colistin | Putative AMP exporter | - |

## Discussion

*S. epidermidis* is a coagulase-negative and Gram-positive staphylococcus that is part of the normal mucosa and skin microflora in humans and other mammals [2]. It is the second leading cause of nosocomial infections [35]. Although it is a saprophyte, opportunistic pathogen with plenty of antibiotic resistance and virulence factors [36], this natural skin colonizer plays a critical role in balancing the epithelial microflora [1, 37]. As an innocuous commensal microorganism, for a long time *S. epidermidis* has been seen as an avirulent species. With the accumulation of genomic sequences, we can now further explore the genetic mechanisms of environmental adaptability of *S. epidermidis*, the evolution process during the outbreak, and the molecular biomarkers for clinic diagnosis [1, 38].

In our current pan-genome analysis, *S. epidermidis* had a relatively compact genome with a size of about 2.5 Mb, and yet almost 20% of this genome was in flux, exchanging with a large pool of various genes. These findings were similar to what had been reported by Conlan and colleagues [29]. The significant number of genes involved in mobilome make horizontal gene transfer easier between the *Staphylococcus* stains and lead to the increase of the “open” pan-genome [39]. Besides, mobile genetic elements, such as SCC*mec*, ACME and plasmids, make the genome structure more unpredictable [40]. High-resolution phylogenetic tree constructed from genome-wide SNPs reveal important details not seen by traditional multi-locus sequence typing (MLST) or single gene marker (16S rDNA). From the phylogenetic tree, we found the ST2 isolates had an extremely short evolutionary distance from each other. The genetic markers *mec*A and *ica*A, which are used to predict the antimicrobial resistance and biofilm phenotypes, have been shown to be more common in hospital isolates than in non-hospital isolates; however, these markers have much less power to distinguish infection isolates from commensally available isolates that contaminate clinical specimens [41]. According to the enrichment analysis, we found it was impossible to distinguish the strains of blood from that of skin, both of which had a similar lifestyle and genetic background. However, it is possible to identify the strains from other habitats with biomarkers such as *ica*ABCD and *cap*ABCD. Whole genome sequencing has been proved to be a more powerful routine diagnostic tool than the traditional MLST or RT-PCR because it can rapidly identify the infection source and antibiotic resistance in an affordable manner [42, 43]. As more genetic data of *S. epidermidis* have been available and new machine learning algorithm is developed [41], WGS may help to predict the infection isolation sources and antibiotic resistance in a quicker and more accurate manner.

*S. epidermidis* has very complicated relationship with human and other bacteria. Antimicrobial peptides (AMPs) play an important role in providing immunity to bacterial colonization on human epithelia. Recent research has shown that staphylococci have multiple systems to combat AMP activity, including AMP sensor that can regulate the expressions of genes involved in AMP resistance depending on the presence of AMPs [7]. We analyzed the distribution of gene involved in AMP resistance and found significant enrichment in blood and skin and variable in different strains, which may be the consequence of coevolution of human’s immune system. On the other side, *S. epidermidis* strains also can inhibit the growth of other bacterium to be dominant species on the skin surface. Serine protease Esp, which is secreted by *S. epidermidis*, has been found to be able to inhibit the biofilm formation of *S. aureus* and destroy pre-existing *S. aureus* biofilms [6]. Other mechanisms are also involved in fighting against pathogens and maintaining homeostasis [44, 45]. On the other hand, *S. epidermidis* was found to be a reservoir of antibiotic resistance, with its virulence determinants shared with other more pathogenic species such as *S. aureus*, as demonstrated in previous studies [29]. In particular, SCC*mec*, ACME elements conferring β-lactam resistance, and other genes are transferred frequently between *Staphylococcus* strains, enabling rapid evolution and adaptation against antibiotic selection pressure and provide additional competitive advantage. For instance, type III of SCC*mec* carries a phenol soluble modulin *psm*-*mec*, which may affect the virulence of *S. aureus* [40].

In conclusion, our current study provides information on the molecular characteristics of *S. epidermidis* strains isolated from different environments from all over the world. From a genomics perspective, the pan-genome analysis of the *S. epidermidis* reveals a high level of diversity among the generic and species-specific genes and the potential supply routes for enhanced versatility via inter- and intra-species horizontal gene transfer. Frequent horizontal gene transfer enables the *Staphylococcus* to adapt to complex environments (e.g., high-level antibiotic), and it may continue to be the dominant genus over the next few years. The understanding of the mechanisms of gene transfer helps us to better prevent the emergence of epidemic pan-drug resistant *S. epidermidis* strains.

## List of abbreviations

SCC*mec*: Staphylococcal chromosome cassette *mec*
AMPs: Antimicrobial peptides
SNPs: Single-nucleotide polymorphisms
COGs: Clusters of orthologous groups
ST: Sequence type
AMR: Antimicrobial resistance

## Acknowledgments

This work was supported by the National Key Research and Development Program of China (Grant 2018YFC2000500), CAMS Innovation Fund for Medical Sciences (2018-I2M-1-002), the Applied Research Program of Capital Clinical Features (Grant Z18110001718172), the Key Research Program for Health Care in China (Grant W2016ZD01).

The authors declare that they have no competing interests.

## Additional files

Table S1 Basic information of all strains analyzed used in this study

Table S2 Result of COG enrichment analysis across all strains

Table S3 Result of KEGG enrichment analysis across all strains

Table S4 Cluster of orthologous proteins produced by PanOCT

Table S5 Enrichment analysis about antibiotic resistances from different sources

Table S6 Enrichment analysis about genes related in biofilm formation

